# Assessment of the cardiovascular adverse effects of drug-drug interactions through a combined analysis of spontaneous reports and predicted drug-target interactions

**DOI:** 10.1101/543918

**Authors:** Sergey Ivanov, Alexey Lagunin, Dmitry Filimonov, Vladimir Poroikov

**Affiliations:** Department of Bioinformatics, Institute of Biomedical Chemistry, Moscow, Russia; Medico-biological Faculty, Pirogov Russian National Research Medical University, Moscow, Russia

**Author notes:** Corresponding author: (SI).

## Abstract

Adverse drug effects (ADEs) are one of the leading causes of death in developed countries and are the main reason for drug recalls from the market, whereas the ADEs that are associated with action on the cardiovascular system are the most dangerous and widespread. The treatment of human diseases often requires the intake of several drugs, which can lead to undesirable drug-drug interactions (DDIs), thus causing an increase in the frequency and severity of ADEs. An evaluation of DDI-induced ADEs is a nontrivial task and requires numerous experimental and clinical studies. Therefore, we developed a computational approach to assess the cardiovascular ADEs of DDIs.

This approach is based on the combined analysis of spontaneous reports (SRs) and predicted drug-target interactions to estimate the five cardiovascular ADEs that are induced by DDIs, namely, myocardial infarction, ischemic stroke, ventricular tachycardia, cardiac failure, and arterial hypertension.

We applied a method based on least absolute shrinkage and selection operator (LASSO) logistic regression to SRs for the identification of interacting pairs of drugs causing corresponding ADEs, as well as noninteracting pairs of drugs. As a result, five datasets containing, on average, 3100 ADE-causing and non-ADE-causing drug pairs were created. The obtained data, along with information on the interaction of drugs with 1553 human targets predicted by PASS Targets software, were used to create five classification models using the Random Forest method. The average area under the ROC curve of the obtained models, sensitivity, specificity and balanced accuracy were 0.838, 0.764, 0.754 and 0.759, respectively.

The predicted drug targets were also used to hypothesize the potential mechanisms of DDI-induced ventricular tachycardia for the top-scoring drug pairs.

The created five classification models can be used for the identification of drug combinations that are potentially the most or least dangerous for the cardiovascular system.

**Author summary:** Assessment of adverse drug effects as well as the influence of drug-drug interactions on their manifestation is a nontrivial task that requires numerous experimental and clinical studies. We developed a computational approach for the prediction of adverse effects that are induced by drug-drug interactions, which are based on a combined analysis of spontaneous reports and predicted drug-target interactions. Importantly, the approach requires only structural formulas to predict adverse effects, and, therefore, may be applied for new, insufficiently studied drugs. We applied the approach to predict five of the most important cardiovascular adverse effects, because they are the most dangerous and widespread. These effects are myocardial infarction, ischemic stroke, ventricular tachycardia, arterial hypertension and cardiac failure. The accuracies of predictive models were relatively high, in the range of 73-81%; therefore, we performed a prediction of the five cardiovascular adverse effects for the large number of drug pairs and revealed the combinations that are the most dangerous for the cardiovascular system. We consider that the developed approach can be used for the identification of pairwise drug combinations that are potentially the most or least dangerous for the cardiovascular system.

## Introduction

Adverse drug effects (ADEs) are one of the top 10 causes of death in developed countries, are one of the main reasons for stopping the development of new drug-candidates and are the main reason for drug recalls from the market [1, 2]. Cardiovascular effects are some of the most serious ADEs that may lead to hospitalization or death, and, at the same time, are widespread [1]. The ADE profile of a particular drug-candidate is usually investigated during standard preclinical animal tests and clinical trials according to the GLP and GCP requirements. However, many rare, but serious, ADEs cannot be revealed by these studies, because of interspecies differences, the limited number of patients or animals and the duration of studies; thus, additional *in vitro* and *in silico* methods for the detection of serious ADEs are currently being developed [3–8]. These methods are based on the determination of the relationships between several chemical and biological features of drugs and their ADEs. Among these features are molecular descriptors, known and predicted drug targets, gene expression changes induced by drugs, phenotypic features such as perturbed pathways, or known ADEs. The relationships between these features and ADEs are usually established using various machine learning methods and network-based approaches. It is accepted that the interaction with human proteins is the most common cause of ADEs; therefore, known and predicted human targets are the most common type of drug features that are used in corresponding studies. Many of the developed methods require knowledge of only the structural formula of a drug-candidate to predict its potential ADEs; therefore, they can be used at the earliest stages of drug development, which may sufficiently increase their effectiveness [3, 4, 8].

In real clinical practice, the treatment of human diseases often requires the administration of several drugs, which can lead to drug-drug interactions (DDIs), thus causing an increase in the frequency and severity of ADEs [9]. An evaluation of the effect of DDIs on the manifestation of ADEs is a nontrivial task and requires numerous preclinical and clinical studies. To solve this problem various computational approaches for the prediction of DDIs were developed [10–22]. Most of these approaches are based on the calculation of similarities between the profiles of various chemical and biological features of two drugs. These similarities can be calculated based on molecular fingerprints, drug targets, their amino acid sequences, pathways and Gene Ontology (http://www.geneontology.org/) annotations, the Anatomical Therapeutic Chemical (ATC) Classification terms (https://www.whocc.no/atc_ddd_index/), as well as known ADEs of individual drugs [10, 12, 13, 15–17, 18, 20, 22]. The Tanimoto coefficient is the most common similarity that is measure in these studies; however, more complicated measures can be used, e.g., several approaches were developed to calculate the proximity of the protein targets of two drugs in a protein-protein interaction network [12, 17]. Similarity measures based on the profiles of different features can be integrated into single interaction scores that allow drug pairs to be ranked according to their potential ability to interact with each other. To estimate the parameters of such integration and validation of obtained results, information about known DDIs was used. Such data can be obtained from various public databases, including DrugBank (https://www.drugbank.ca/) and Drugs.com (https://www.drugs.com/). For example, Cheng F. with colleagues [13] used several machine learning methods with drug phenotypic, therapeutic, chemical and genomic similarities used as features to predict DDIs. The classifiers were trained on the set of known DDIs from the DrugBank database and the same number of randomly chosen drug pairs as the negative examples. The best result with the area under the ROC-curve (AUC) 0.67 was achieved using a support vector machine with a Gaussian radial basis function kernel. In addition to approaches that are based on similarities, some other methods were developed [14, 19]. Zakharov A.V. with colleagues [19] used separate training sets of pairwise drug combinations for each of four isoforms of cytochromes P450, which are examples of known DDIs. The corresponding information was obtained from the literature. Drug pairs were represented as mixtures of compounds in ratio 1:1, and several types of molecular descriptors were generated for them. The prediction models were generated by using the radial basis function self-consistent regression and a Random Forest. The balanced accuracies that were obtained from the cross-validation procedure varied from 0.72 to 0.79, depending on the dataset [19]. Luo H., with colleagues, used the sums and differences of the docking scores for 611 human proteins to describe 6328 drug pairs, which represented known DDIs from the DrugBank database, and the same number of drug pairs was randomly chosen as a negative example. A predictive model was created based on l2-regularized logistic regressions to obtain their values. The obtained accuracy, sensitivity and specificity that were calculated based on the 10-fold cross-validation procedure were 0.804, 0.847 and 0.772, respectively [14].

Despite the significant progress in predicting DDIs, all of these methods allow for estimating only the fact of interaction, but not the resulting ADEs, whereas such information is important to assess the clinical significance of DDIs. The main problem is the absence of known data for most of the DDI-induced ADEs. The major source of data on ADEs of individual drugs is drug labels [23]; however, they usually contain very few data on ADEs that are induced by DDIs. Nevertheless, the corresponding information can be obtained through the analysis of spontaneous reports (SRs) which are received by regulatory agencies from healthcare professionals and patients. Each SR contains information about all drugs that are prescribed to a patient, as well as information about developed ADEs. An analysis of large sets of SRs allows for relationships between certain ADEs and individual drugs [24–29], or drug combinations [30–35], to be revealed. The datasets of individual drugs with information about ADEs obtained by an analysis of SRs were earlier successfully used for the creation of predictive models that are based on structure-activity relationships [27, 29]. The corresponding information on ADEs that is induced by pairwise drug combinations may also potentially be used for this purpose.

We developed a computational approach for the assessment of cardiovascular ADEs of DDIs. The approach is based on a combined analysis of SRs and predicted drug-target interactions (DTIs) and allows for the prediction of five cardiovascular ADEs of DDIs: myocardial infarction, ischemic stroke, ventricular tachycardia, arterial hypertension and cardiac failure, with balanced accuracies from 0.73 to 0.81. Unlike most of the other methods, our approach requires only structural formulas to predict cardiovascular adverse effects for any pair of drugs, and, therefore, may be applied for new, drug-like compounds that have not yet been studied. The developed approach can be used for the identification of pairwise drug combinations that are potentially the most or least dangerous for the cardiovascular system.

## Results and discussion

### General description of the approach

We developed a new computational approach for the assessment of cardiovascular ADEs of DDIs through a combined analysis of SRs and predicted DTIs (Fig 1).

**Fig 1.**
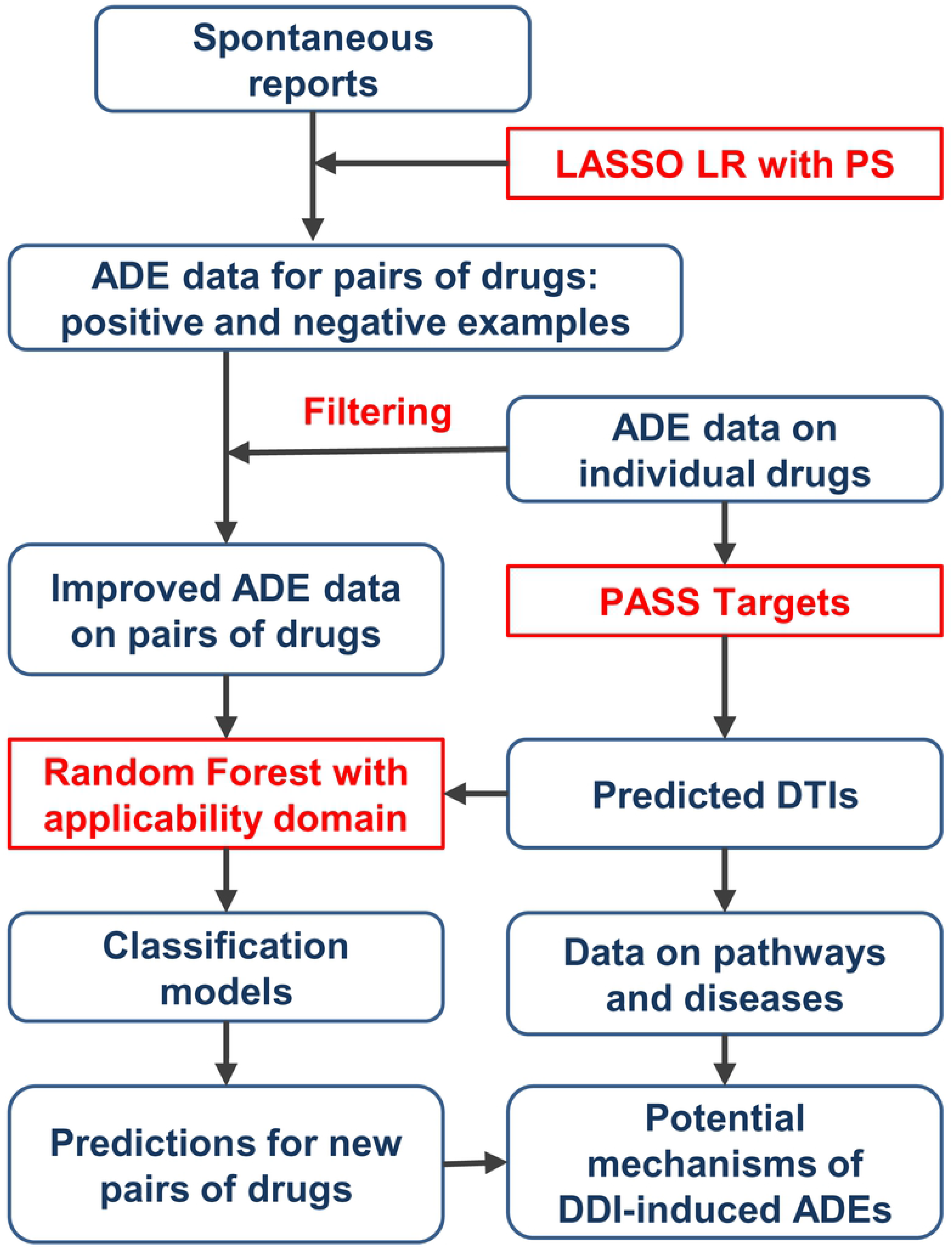
The scheme of a developed computational approach for the assessment of cardiovascular ADEs of DDIs. LASSO LR – least absolute shrinkage and selection operator (LASSO) logistic regression, PS – propensity scores (see Material and Methods).

The approach is based on two main steps: creation of datasets on cardiovascular DDI-induced ADEs containing drug pairs that cause or do not cause ADEs, and the creation of classification models for each dataset based on predicted drug targets as descriptors. The creation of datasets is based on the analysis of SRs from the standardized version of publicly available parts of the FDA database [36]. The analysis was performed using least absolute shrinkage and selection operator (LASSO) logistic regression with the addition of propensity scores as independent variables [35] (see Materials and Methods for details), which allows for the identification of drug pairs that cause or do not cause cardiovascular ADEs – positive and negative examples. Each “positive” drug pair represents a synergistic or additive effect of DDI on the development of ADEs. This method takes into account the confounding effects of other drugs and risk factors on the manifestation of ADEs and, thus, allows for datasets with lower numbers of false positives to be obtained. To further improve the quality of datasets, information about the ADEs of individual drugs [37] was used to filter out potentially false positive and false negative examples (see Materials and Methods).

At the second step of the approach, a PASS Targets software [38] was used to predict interactions of individual drugs that were from obtained datasets with 1553 human protein targets. The sums and absolute values of the differences in the probability estimates of interaction with targets were used as descriptors for drug pairs. The classification models were built using Random Forest along with a method that allows for the applicability domain to be determined. The accuracy of prediction is estimated using a 5-fold cross-validation procedure (see Materials and Methods). To demonstrate the practical benefit of the obtained models, predictions of ADEs for a large amount of drug pairs were performed. The analysis of the biological role of predicted protein targets for the top predicted drug pairs that potentially cause ADEs allows for proposing the potential mechanisms of corresponding DDIs.

### Creation of datasets

At the first step of the proposed approach, we created five datasets of drug pairs that cause and do not cause five cardiovascular ADEs through the analysis of SRs (see Materials and Methods), namely, ventricular tachycardia, myocardial infarction, ischemic stroke, arterial hypertension and cardiac failure (see Table S1). Each positive drug pair represents an example of a synergistic or additive DDI that causes a corresponding ADE. The datasets contain, on average, more than 3100 drug pairs belonging to 335 individual drugs and 166 ATC terms of the fourth level (Table 1).

**Table 1.** Characteristics of created datasets on DDI-induced ADEs.

|  | Positive pairs | Negative pairs | Number of drugs | Number of ATC classes |
| --- | --- | --- | --- | --- |
| Ventricular tachycardia | 933 | 2912 | 376 | 181 |
| Myocardial infarction | 2479 | 1279 | 352 | 168 |
| Ischemic stroke | 838 | 2101 | 331 | 169 |
| Arterial hypertension | 549 | 1029 | 273 | 146 |
| Cardiac failure | 1350 | 2108 | 343 | 166 |

We performed the following analysis to estimate whether the obtained datasets contain information reflecting DDI-induced ADEs or datasets containing information that is similar to random. One may suggest that the obtained datasets contain more positive drug pairs where both drugs cause the ADE when administered separately (both-ADE-causing pair) than expected by chance. Indeed, the induction of ADE by DDI is more probable when both drugs may cause a particular ADE. Similarly, one may suggest that the obtained datasets contain more positive drug pairs, where only one of the two drugs causes the ADE (one-ADE-causing pair), compared to the positive drug pairs, where neither of the two drugs cause ADE (none-ADE-causing pair), than expected by chance. We compared percentages of both-, one- and none-ADE-causing drug pairs among the positive pairs of datasets to the corresponding percentages for background datasets (Fig 2). The background datasets contained all pairwise drug combinations where information about corresponding ADEs of individual drugs was available, and both drugs were simultaneously mentioned in at least 100 SRs. The obtained result indicates that the positive drug pairs from our datasets were significantly enriched with both- and one-ADE-causing pairs compared to the background.

**Fig 2.**
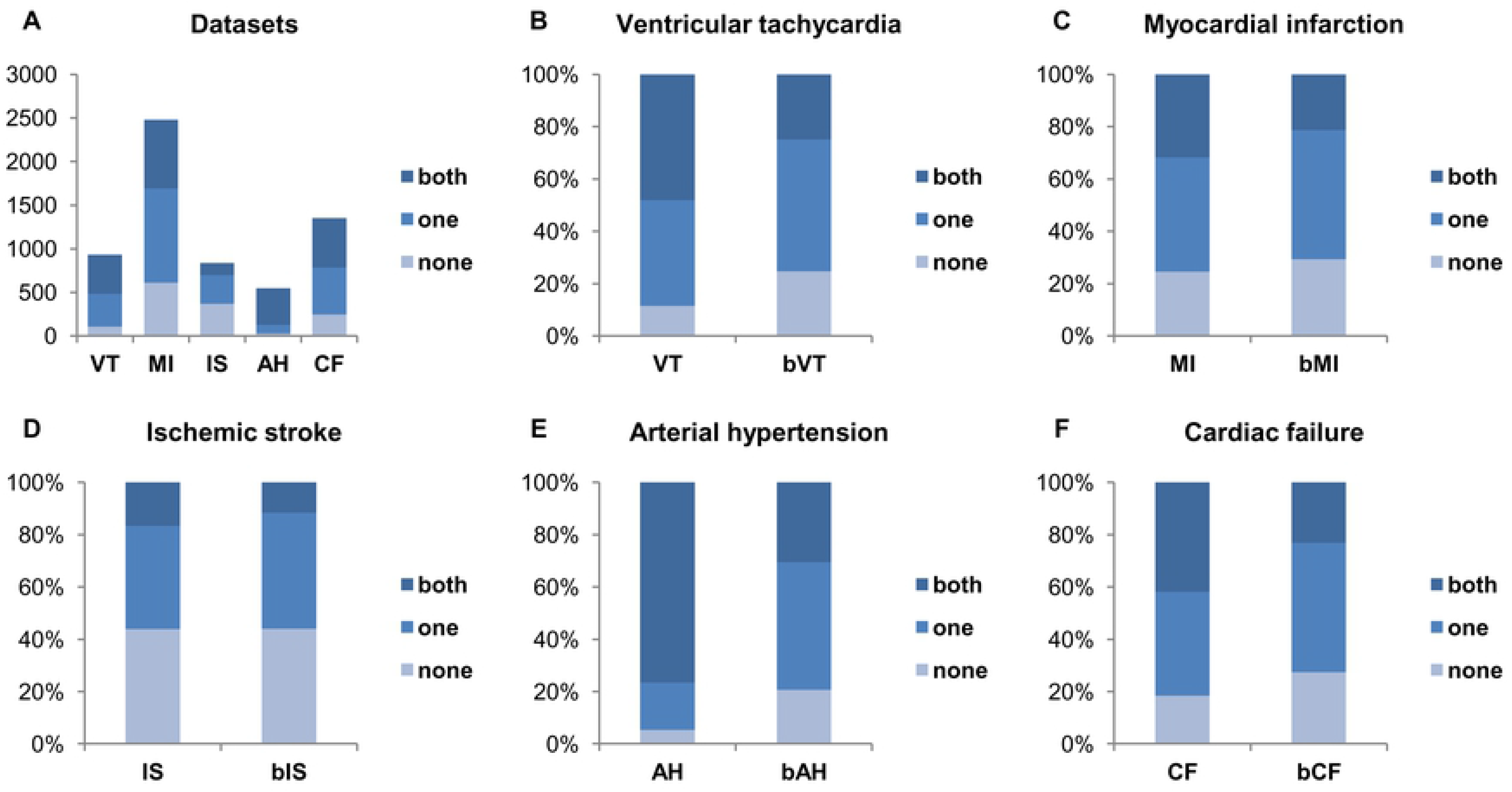
Percentages of both-, one- and none-ADE-causing drug pairs among positive pairs of obtained and background datasets. VT – ventricular tachycardia, MI – myocardial infarction, IS – ischemic stroke, AH – arterial hypertension, CF – cardiac failure; VT, MI, IS, AH, and CF are positive drug pairs from the created datasets; bVT, bMI, bIS, bAH, and bCF are drug pairs from background datasets.

The statistical significance of enrichment was estimated using the chi-squared test. Enrichments for all five ADEs were statistically significant with the highest p-value 0.000035 for ST dataset.

As a result, the created datasets are relevant, representative and can be used for further analysis.

### Prediction of DDI-induced cardiovascular ADEs based on drug-target interactions

We used Random Forest to create classification models and the local (Tree) approach to determine their applicability domain [39]. The models were created based on sums and absolute values of differences of probability estimates of interaction with 1553 human protein targets that had been calculated for individual drugs by PASS Targets software [38]. The accuracy estimates were obtained by a 5-fold cross-validation procedure with use of the “compound out” approach [40] (see Materials and Methods for details). The obtained average values of AUC, sensitivity, specificity and balanced accuracy were 0.838, 0.764, 0.754 and 0.759, respectively, whereas 95.7% of the drug pairs were in the applicability domain of the models (Table 2).

**Table 2.** Prediction accuracy for five cardiovascular DDI-induced ADEs based on 5-fold cross-validation procedure.

|  | AUC | Sensitivity | Specificity | Balanced accuracy | In applicability domain |
| --- | --- | --- | --- | --- | --- |
| Ventricular tachycardia | 0.807 | 0.743 | 0.718 | 0.731 | 96.1% |
| Myocardial infarction | 0.856 | 0.794 | 0.763 | 0.778 | 95.3% |
| Ischemic stroke | 0.808 | 0.734 | 0.724 | 0.729 | 95.6% |
| Arterial hypertension | 0.892 | 0.789 | 0.832 | 0.810 | 95.5% |
| Cardiac failure | 0.824 | 0.761 | 0.734 | 0.747 | 96.1% |

The obtained relatively high accuracies allow for the application of the created models to solve practical tasks, e.g., to perform a search of new pairwise combinations of drugs that potentially interact and cause cardiovascular ADEs.

### Prediction of DDI-induced ADEs for the new drug pairs

The created datasets contain from hundreds to thousands of drug pairs that cause cardiovascular ADEs depending on the effect; however, the number of possible pairwise drug combinations is much higher. To investigate the practical benefit of the created classification models, we performed a prediction of the DDIs-induced ADEs for all of the possible drug pairs that were generated from individual drugs with known data on five cardiovascular ADEs [37]. Five large datasets were generated with more than 230000 drug pairs on average, and 190000 pairs (84%) of them were in the applicability domain of the models (see Table 3). Surprisingly, nearly half of the drug pairs in the datasets were predicted to cause corresponding DDI-induced ADEs. The average number of positive drug pairs in the training sets (see Table 1) is almost approximately 40%. Moreover, according to the datasets of individual drugs with information on five cardiovascular ADEs taken from our previous work [37], nearly 40% of the single drugs also cause ADE. This is not much less than the percentages of predicted ADE-causing drug pairs (Table 3).

**Table 3.** Numbers of drug pairs with predicted ADEs.

|  | Number of pairs | Number of pairs with ADE | Pairs with ADE, % | In applicability domain, % |
| --- | --- | --- | --- | --- |
| Ventricular tachycardia | 232480 | 157393 | 52.8 | 84.9 |
| Myocardial infarction | 195933 | 101063 | 51.6 | 83.3 |
| Ischemic stroke | 189275 | 98006 | 51.8 | 84.5 |
| Arterial hypertension | 161873 | 57457 | 35.5 | 86.2 |
| Cardiac failure | 192326 | 108064 | 56.2 | 82.8 |

The high percentages of known and predicted ADE-causing drug pairs may be explained by the fact that most of them, such as individual drugs, may cause ADEs only in a small percentage of patients. Therefore, the clinical significance of DDI in relation to the cardiovascular system may be estimated based on only the number of predicted ADEs. We chose 63212 drug pairs that are in all five of the large datasets and are in the applicability domain of all five models and found that only 4707 drug pairs (7.4%) potentially cause all five cardiovascular ADEs (Fig 3). The potentially most dangerous drug combinations are listed in Table S2.

**Fig 3.**
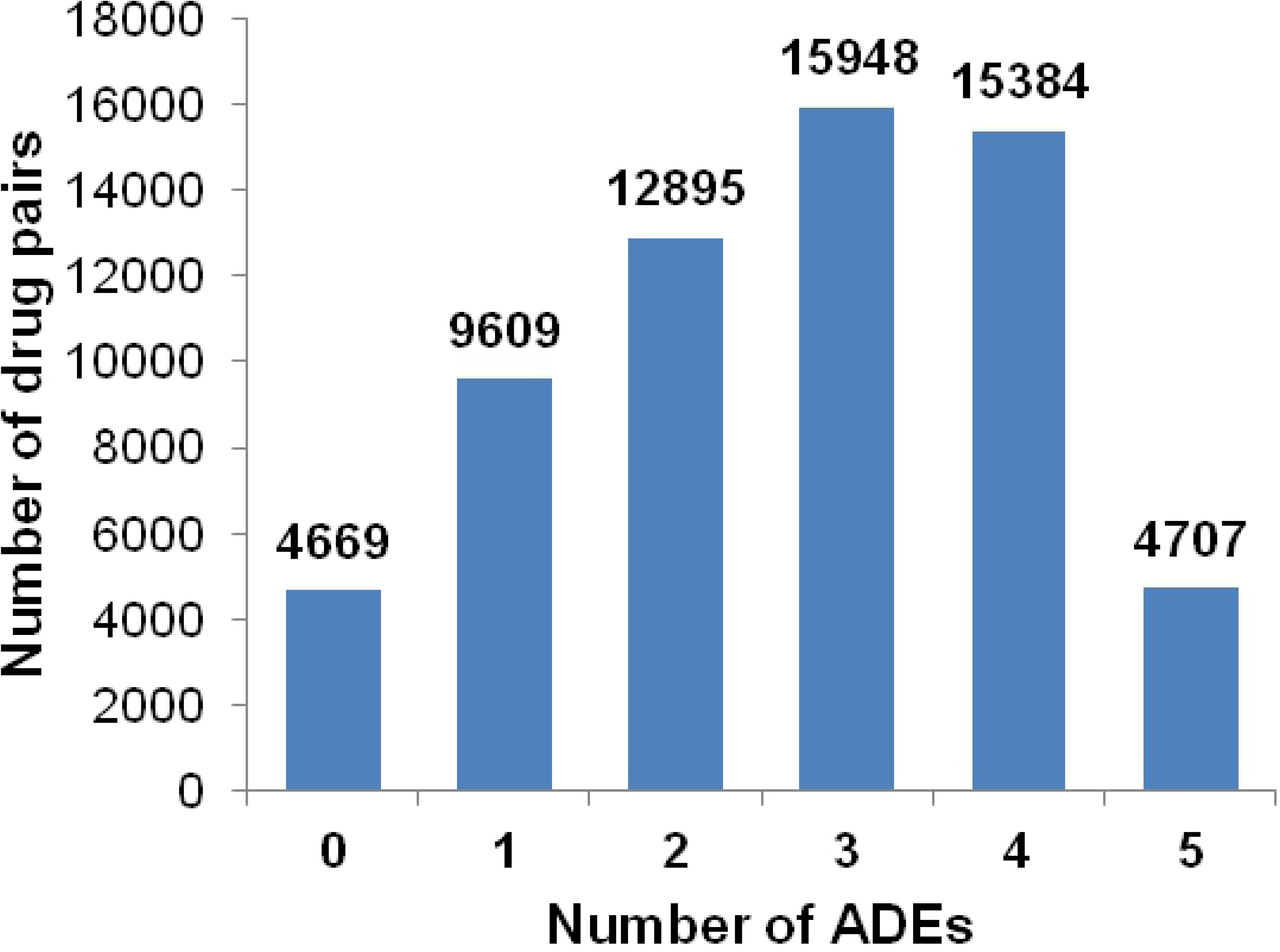
Number of cardiovascular ADEs predicted for a large dataset of drug pairs. Zero (“0”) means that none of five ADEs were predicted for the pairwise drug combination.

To estimate the relevance of the predicted ADEs, we performed an analysis of the distribution of both-, one- and none-ADE-causing drug pairs among combinations that were predicted to be positives and negatives in relation to the corresponding ADEs (Fig 4).

**Fig 4.**
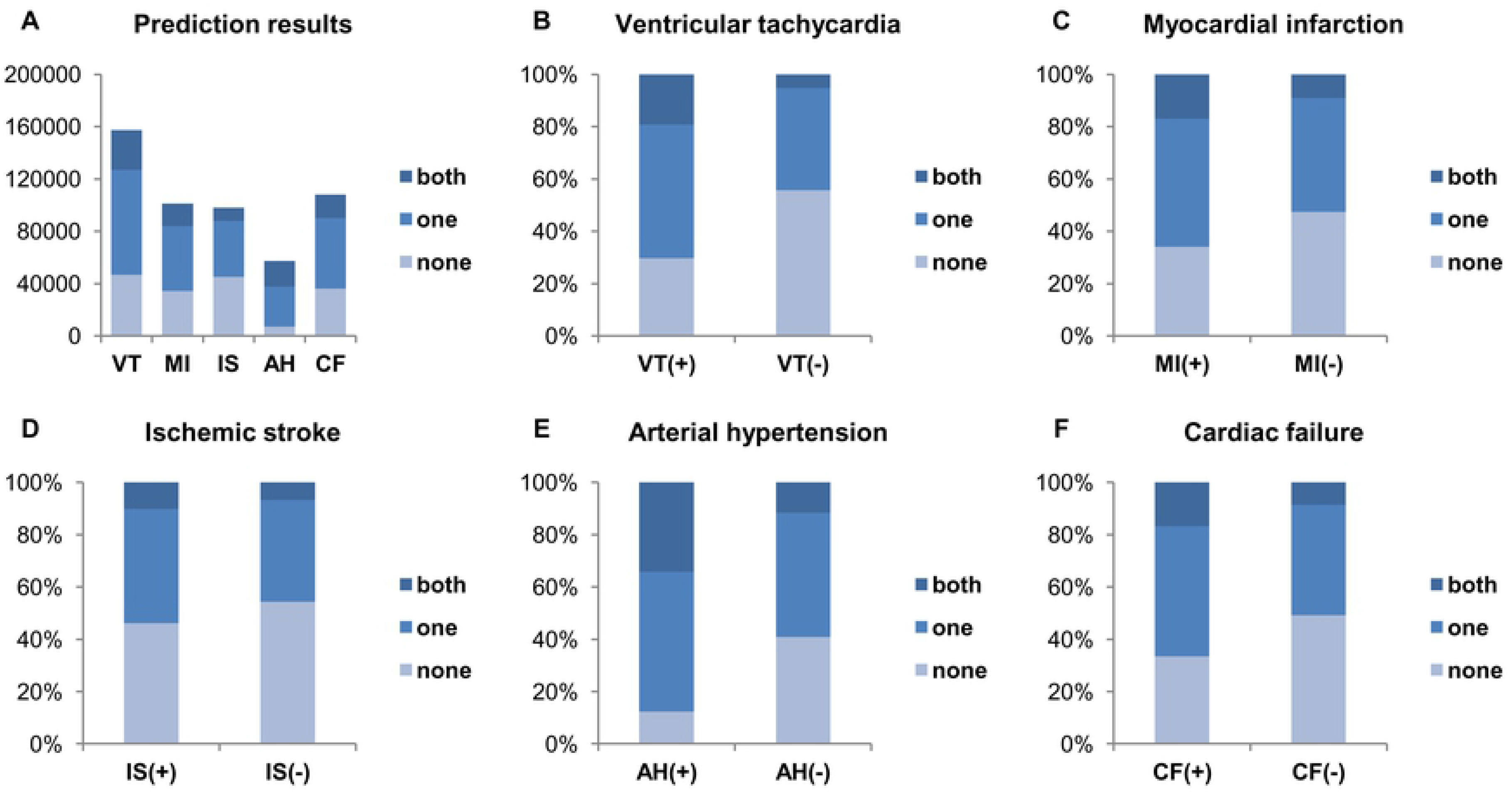
Percentages of both-, one- and none-ADE-causing drug pairs among predicted positive and negative pairs of large datasets. VT(+), MI(+), IS(+), AH(+), and CF(+) are predicted to be positive drug pairs for ventricular tachycardia, myocardial infarction, ischemic stroke, arterial hypertension and cardiac failure; VT(-), MI(-), IS(-), AH(-), and CF(-) are predicted to be negative drug pairs for the same ADEs.

The observed distribution is similar to that shown in Fig 2. We found that drug pairs that potentially cause ADE contain higher percentages of both- and one-ADE-causing pairs compared to the drug pairs that potentially do not cause ADE. It means that, generally, the prediction results are relevant.

The DrugBank database contains some data on known DDIs that lead to ventricular tachycardia (or prolongation of the QT interval on an electrocardiogram) and arterial hypertension. We selected corresponding drug pairs that intersect with the large created datasets and are in the applicability domain of classification models (Table 4).

**Table 4.** Prediction accuracy on positive drug pairs from the DrugBank for ventricular tachycardia and arterial hypertension.

|  | N of drug pairs | AUC |
| --- | --- | --- |
| Ventricular tachycardia | 3264 | 0.746 |
| Arterial hypertension | 112 | 0.774 |

We compared the predicted probability estimates of these pairs with all other pairs in large datasets. The observed AUC values indicate that the positive pairs from DrugBank are usually the top ranking among all pairs in the datasets (Table 4).

The results of these analyses and the results of 5-fold cross-validation (the average area under the ROC curve, sensitivity, specificity and balanced accuracy were 0.838, 0.764, 0.754 and 0.759, respectively; see Table 2) indicate that the accuracy of the prediction of DDI-induced cardiovascular ADEs is relatively high and that the created models can be applied in the search for new pairwise combinations of drugs that are the most or the least dangerous for the cardiovascular system. Because DTIs are needed for the creation of models that were predicted by PASS Targets software based on structures of drugs, the developed models can be used for any drug-like compounds, including those for which only structural formulas are known. For example, they can be used to predict DDI-induced ADEs for drug candidates on the stage of clinical trials.

### Assessment of the potential mechanisms of DDI-induced ADEs

Since DDI-induced ADEs are effectively estimated by using data on predicted DTIs, the corresponding information on drug targets may also be used to reveal the potential mechanisms of cardiovascular ADEs and to influence DDIs in their manifestation.

We performed a corresponding analysis for the top 5 none-VT-causing drug pairs from the large dataset with the highest probability scores for ventricular tachycardia (VT) (Table 5). The drugs from these pairs cannot cause ventricular tachycardia when administered separately; however, the drugs possibly cause VT when they are administered together.

**Table 5:** Potential mechanisms of DDI-induced ventricular tachycardia (VT) for the top 5 scored none-VT-causing drug pairs. The bold and underlined gene names mean known, experimentally confirmed drug targets from DrugBank and ChEMBL databases. Symbols ↑ and ↓ mean up- and down-regulation of the protein function by the drug.

| Drug pairs | Common cytochromes P450 | Known and predicted drug targets associated with ventricular tachycardia |
| --- | --- | --- |
| Dapsone-Emtricitabine | – | Dapsone: SGK3. Emtricitabine: PRKAA2, ULK1 |
| Cortisone Acetate-Dapsone | <u>CYP3A4</u> | Cortisone Acetate: **NR3C1**↑. Dapsone: SGK3 |
| Eszopiclone-Chlorphenamine | <u>CYP3A4</u> | Eszopiclone: **TSPO**↑, CAMKK1, ULK1. Chlorphenamine: **HRH1**↓, **SLC6A2**↓, **HTR2B**, HRH2, KCNH2, CALM |
| Dapsone-Dolutegravir | <u>CYP3A4</u> | Dapsone: SGK3 |
| Voriconazole-Dapsone | <u>CYP3A4</u>, <u>CYP3A5</u>, <u>CYP3A7</u>, <u>CYP2C9</u>, <u>CYP2C19</u> | Voriconazole: HSP90AA1. Dapsone: SGK3 |

We found that the DDIs for these drug pairs may occur at both levels of pharmacokinetics and pharmacodynamics. First, the drugs from four of five pairs are metabolized by the same cytochromes P450. Second, corresponding drugs potentially interact with protein targets to influence the action potential of cardiac cells. These targets, either known or predicted, are shown in Table 5. It is important that only chlorphenamine was predicted to interact with the HERG (KCNH2) potassium channel, which is a well-known protein that is associated with ventricular tachycardia [5]. However, this and other drugs from selected pairs that are known to or are predicted to interact with human proteins form compact fragments of the regulatory network (Fig 5) and indirectly change the action potential. Such changes may form a basis for the induction of ventricular tachycardia in predisposed patients.

**Fig 5.**
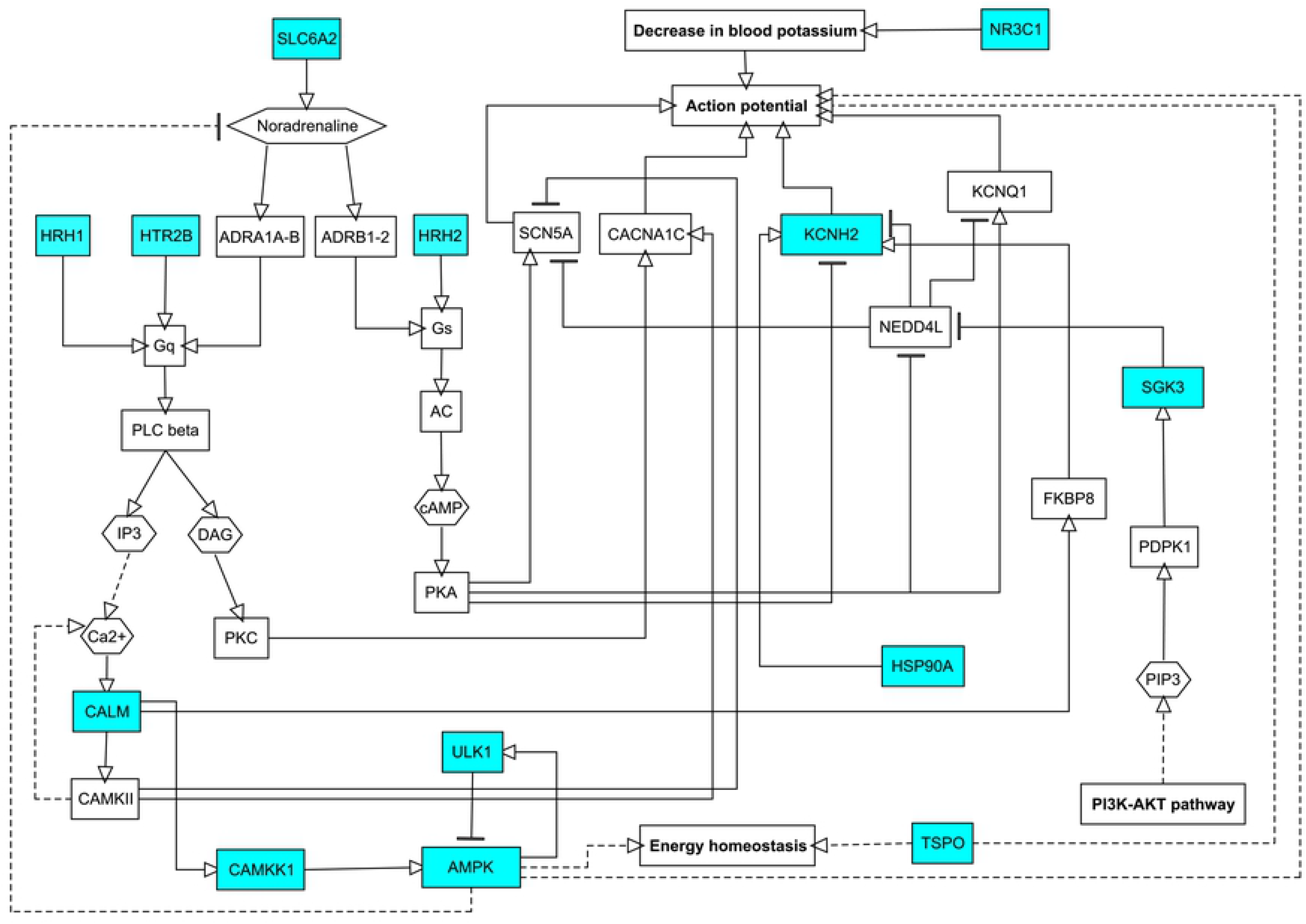
Influence of known and predicted protein targets of the top 5 scored none-VT-causing drug pairs on the action potential in the heart. VT – ventricular tachycardia. Cyan nodes represent known and predicted protein targets of drugs from selected pairs, and white nodes represent intermediate proteins in the regulatory network. Solid edges represent direct interactions, and dashed edges represent indirect interactions. The figure was created based on data from KEGG pathways (https://www.genome.jp/kegg/pathway.html) and from corresponding information in the literature.

## Materials and Methods

### Assessment of DDI-induced ADEs through the analysis of SRs

In our study, we used the AEOLUS database [36] as a source of SRs. AEOLUS is a curated version of publicly available parts of the FDA database of SRs (https://www.fda.gov/Drugs/GuidanceComplianceRegulatoryInformation/Surveillance/Adverse_DrugEffects/default.htm), where the names of ADEs, drugs and indications are standardized. We selected only those SRs that contain description of drugs, ADEs and drug indications, because all of these types of data are required for further analysis. A total of 4028051 SRs were selected. The ADEs and indications in the database were described by the preferred terms (PTs) of the MedDRA dictionary (https://www.meddra.org/). Since some PTs may describe pathologies that are related to the same or similar ADEs, we selected the main PTs, which exactly match the investigated ADEs and support PTs, which are conditions that are similar to or are indirectly related to ADEs. The main and supporting PTs for five investigated cardiovascular ADEs are presented in Table 6.

**Table 6.** Main and supporting PTs for five investigated cardiovascular ADEs.

| Main PTs | Supporting PTs |
| --- | --- |
| Torsade de Pointes<br>Ventricular Tachycardia | Electrocardiogram QT Prolonged<br>Electrocardiogram QT Corrected Interval Prolonged<br>Ventricular Arrhythmia |
| Acute Myocardial Infarction<br>Acute Coronary Syndrome<br>Myocardial Infarction | Angina Pectoris<br>Angina Unstable<br>Arteriosclerosis Coronary Artery<br>Arteriospasm Coronary<br>Coronary Artery Disease<br>Coronary Artery Occlusion<br>Coronary Artery Stenosis<br>Coronary Artery Thrombosis<br>Myocardial Ischemia |
| Hypertension<br>Hypertensive Crisis | Blood Pressure Increased<br>Blood Pressure Systolic Increased<br>Blood Pressure Diastolic Increased |
| Cerebrovascular Accident<br>Cerebral Infarction<br>Ischemic Stroke | Cerebral Ischemia<br>Transient Ischemic Attack |
| Cardiac Failure Acute<br>Cardiac Failure Congestive<br>Cardiac Failure<br>Cardiogenic Shock<br>Cardiopulmonary Failure<br>Left Ventricular Failure<br>Right Ventricular Failure | — |

At the next step, we selected those drugs in the AEOLUS database that have annotations on five investigated cardiovascular ADEs: ventricular tachycardia, myocardial infarction, ischemic stroke, arterial hypertension and cardiac failure. The data on drugs that caused and did not cause five ADEs was obtained from our previous study [37]. The following numbers of drugs were selected: 496 drugs for ventricular tachycardia, 460 drugs for myocardial infarction, 447 drugs for ischemic stroke, 398 drugs for arterial hypertension, and 467 drugs for cardiac failure. The data on the five ADEs of these individual drugs are represented in Table S3.

We selected drug pairs that were formed by these drugs with at least 100 SRs wherein both drugs are mentioned. For each pair of drugs and each PT from Table 6, we performed an analysis which is based on three steps. At the first step, we found which of the drug pairs are associated with selected PTs. At the second step we used LASSO logistical regression [35] to estimate the potential synergistic and additive DDIs that are associated with the drug pairs that were selected in step 1. At this step, noninteracting drug pairs were also determined. At the third step, we integrated the obtained data on different PTs into single ADEs to create datasets with positive and negative examples of DDI-induced ADEs (see Table 1).

#### Step 1. Identification of the association between drug pairs and PTs

A proportional reporting ratio (PRR) was used to determine the drug pairs that are associated with each PT. PRR is calculated as follows:

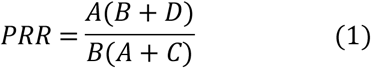

The value A is a number of the SRs where both the drug pair and PT are mentioned; B is a number of SRs where PT is mentioned, but the drug pair is not mentioned; C is a number of SRs where the drug pair and other PTs are mentioned; and D is a number of SRs where the PT and drug pair are not mentioned.

According to previously published criteria [26, 28], we considered a relationship between the drug pair and PT if PRR>2, A>2 and chi-square≥4. The selected associations were used at the next step of analysis.

#### Step 2. Identification of synergistic and additive DDIs

We identified synergistic and additive pairwise DDIs that are associated with each PT by using LASSO logistic regression with propensity scores (PSs). The method is described in detail in the original publication [35].

Briefly, PS is a conditional probability of being exposed to a drug that is calculated for each SR. This probability depends on the patient’s diseases and, indirectly, on co-administered drugs. The PS indirectly reflects the influence of human diseases and co-administered drugs on the development of ADE, and, thus, allows for the filtering of many false positive drug-ADE associations. We calculated the PSs for each drug-SR pair based on the top 100 co-administered drugs and the top 100 most relevant drug indications. The relevance of co-administered drugs and indications of a drug were measured by a phi correlation coefficient.

The final values of the PSs were calculated by using the following logistic regression:

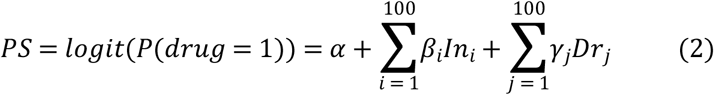

In formula (2), the values In_i_ and Dr_j_ are the indication and co-administered drug with relevance ranks i and j.

Next, we used LASSO logistic regression to estimate the probability of PT for each SR that depends on the presence of two drugs in SR, their possible interaction, and the corresponding PSs as follows:

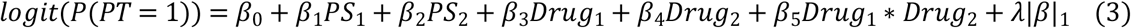

In formula (3), PS_1_ and PS_2_ are PSs for drug_1_ and drug_2_, |β|_1_ is 1_1_ norm of coefficients, and λ is a tuning parameter of regularization. Parameter λ was determined through a 3-fold cross-validation procedure. The potential synergistic and additive DDIs that are associated with PTs were determined based on β_3_, β_4_ and β_5_ coefficients:

- *synergistic DDI* for drug pair-PT association was considered if β_5_ was more than 0;
-*additive DDI* for drug pair-PT association was considered if β_5_ equals 0, β_3_ and β_4_ were more than 0, and drug_1_, drug_2_ have known links to the corresponding ADE in datasets from our previous study [37].
-*absence of DDI* for the drug pair-PT association was considered if either β_3_ or β_4_ were less or equal to 0, and β_5_ was less or equal to 0. Additionally, we considered the absence of DDIs if the corresponding drug pair-PT association was not determined at step 1 (the condition PRR>2, A>2 and chi-square>4 was not true) and the drug pair was mentioned in at least 500 SRs.

#### Step 3. Integration of data on different PTs

To create final datasets with the information on DDI-induced ADEs, we integrated data on the PTs as follows:

- The drug pair was considered to be “positive” according to the corresponding ADE if it was linked to at least two main PTs, or at least to one main and one supporting PT at step 2 of the analysis.
- The drug pair was considered to be “negative” according to the corresponding ADE if it was linked to neither of the PTs that are associated with this ADE. Additionally, we removed from this category those drug pairs in which both drugs are ADE-causing, according to data from our previous study [37], as potentially false negatives.

As a result, datasets for the five cardiovascular ADEs were created (see Results and Discussion, Table 1).

### Prediction of drug-target interactions

Interactions of individual drugs with human proteins were predicted by the PASS Targets software [38]. PASS (Prediction of Activity Spectra for Substances) [41–43] can be used for the prediction of various types of biological activities and is associated with several hundred success stories of its practical application, with experimental confirmation of the prediction results [43, 44]. It uses Multilevel Neighborhoods of Atoms (MNA) descriptors and the Bayesian approach and is available as a desktop program as well as a freely available web service on the Way2Drug platform (http://www.way2drug.com/PASSOnline/) [45]. PASS Targets is a special version of PASS that is based on training data from the ChEMBL database (https://www.ebi.ac.uk/chembl/) and allows for predicting interactions with 1553 human protein targets with an average AUC 0. 97 and a minimal AUC 0.85 [38]. The full list of human targets is presented in Table S4.

PASS provides two estimates of probabilities for each target of a chemical compound: The Pa probability to interact with a target, and the Pi probability to not interact with a target. If a compound has Pa > Pi, it can be considered as interacting with the target. The larger the Pa and Pa-Pi values, the greater the probability of obtaining an activity against a target in the experiment. In this study, we used a threshold Pa>0.3 for the estimation of protein targets of drugs from the top 5 scored non-VT-causing drug pairs (see the last section of the Results and Discussion).

We used sums and absolute values of differences of Pa/(Pa+Pi) values, calculated by PASS for individual drugs, to obtain corresponding values for pairs of drugs. Thus, each drug pair was described by a vector of 3106 values, which were further used as descriptors for the creation of classification models (see below).

### Creation of classification models for DDI-induced cardiovascular ADEs

Classification models for the prediction of five DDI-induced cardiovascular ADEs were created by the r Random Forest method. We used the RandomForest function from “RandomForest” R package (https://cran.r-project.org/web/packages/randomForest/) for this purpose. All arguments of this function were set to default.

The applicability domain of the obtained models was determined by the local (Tree) approach, which was described earlier [39].

The accuracy of created models was determined by a 5-fold cross validation procedure according to the “compound out” approach, wherein each drug pair in the test set must contain at least one drug that is absent in all drug pairs of the training set [40].

## Supporting information

**S1 Table. Datasets with information of DDI-induced cardiovascular ADEs.**

**S2 Table. Drug pairs potentially causing all five cardiovascular ADEs.**

**SA3 Table. Information about cardiovascular ADEs of individual drugs used in the study.**

**S4 Table. The list of human protein targets predicted by PASS Targets software.** Numbers of active compounds in the training set as well as the AUC values that were obtained by leave-one-out cross-validation are given.

